# PGRL1A redox states alleviate photoinhibition in Arabidopsis during step changes in light intensity

**DOI:** 10.1101/2022.06.07.492398

**Authors:** Amit Kumar Chaturvedi, Orly Dym, Robert Fluhr

## Abstract

Non-motile plants have evolved regulatory mechanisms to maintain homeostasis for optimal growth. Responses to environmental changes in light are particularly important not only during the diurnal transition from night to day but also to react to light changes caused by passing clouds or by wind. Thioredoxins rapidly orchestrate redox control during environmental change by modifying cysteine residues. Here, we assign a function to regulatory cysteines of PGRL1A, a constituent of the ferredoxin-dependent cyclic electron flow (Fd-CEF) pathway and show their role in the regulation of proton motive force (PMF) and nonphotochemical quenching (NPQ). During step increase of low light intensity (10-60 μE*m^-2^*s^-1^), the intermolecular disulfide of the PGRL1A 59-kDa complex is reduced transiently within seconds to the 28 kDa form. In contrast, step increases to higher light intensity (60-600 μE*m^-2^*s^-1^) stimulated a stable partially reduced redox state in PGRL1A. Measurements of NPQ, PMF and resultant photosynthetic controls Y(ND) and Y(NA) were found to correlate with the redox state of PGRL1A during step increases in light intensity but not in PGRL1mutant plants *pgrl1ab* or PGRL1A cysteine mutant (*PGRL1A_C1,2A_*). Continuous light regimes did not affect mutant growth; however, fluctuating regimes of light intensity showed significant growth reduction in the mutants. Inhibitors of photosynthesis placed control of the PGRL1A redox state as dependent on the penultimate ferredoxin redox state that fuels reducing equivalents to the large set of chloroplasts thioredoxins. Our results showed that redox state changes in PGRL1A are crucial to the optimization of photosynthesis and are regulated by the photosynthetic electron flux.

## Introduction

In chloroplasts, redox regulation serves to reconfigure plant metabolism in response to rapid fluctuations in light intensity in night-day transitions and during the day (Schürmann and Buchanan, 2008; Allahverdiyeva et al., 2014; Lee et al., 2019). Plants evolved several photoprotective mechanisms that attenuate light capturing reactions by dissipating excess absorbed light as heat through non-photochemical quenching (NPQ) of chlorophyll fluorescence (Pascal et al., 2005; Niyogi and Truong, 2013; Ruban and Ruban, 2018; Murchie and Ruban, 2020). The energy-dependent quenching/feedback de-excitation (qE) is the fastest component of NPQ being modulated within seconds in response to changes in light intensity. The fast and reversible mechanisms that are shown to influence qE include at least 2 mechanisms: modulation of Photosystem II subunit S (PsbS) (Li et al., 2002; Sylak-Glassman et al., 2014) and the xanthophyll cycle. In the latter reaction, the enzyme violaxanthin de-epoxidase (VDE) converts the carotenoid violaxanthin into zeaxanthin by disulfide exchange reactions (Demmig-Adams and Iii, 1996; Jahns and Holzwarth, 2012; Simionato et al., 2015). PsbS acts as a pH sensor to detect lumen acidification and then change the chlorophyll aggregation state (Krishnan-Schmieden et al., 2021; Nicol and Croce, 2021). PsbS-mediated mechanism is considered to be the fastest; reacting to environmental changes within seconds (Li et al., 2002). Both mechanisms have been shown to be regulated by a trans-thylakoid proton gradient (Demmig-Adams and Iii, 1996; Li et al., 2002; Jahns and Holzwarth, 2012; Sylak-Glassman et al., 2014).

Linear electron flow (LEF) is generally limited by the level of its main stromal substrate, oxidized NADP^+^, that is present at the acceptor side of photosystem I (PSI). This is called acceptor side limitation. Hence, to prevent excess flow at the transition period from night to day or during light intensity changes, an alternative pathway, the ferredoxin-dependent cyclic electron flow (Fd-CEF) bypasses NADP^+^ limitations. CEF is thought to have an important photoprotective role and also fulfill the demand of extra ATP to maintain the ATP/NADPH ratio for primary metabolism. (Munekage et al., 2002; Munekage et al., 2004; Johnson, 2005; Shikanai, 2007; Eberhard et al., 2008; Hertle et al., 2013; Yamori and Shikanai, 2016). The CEF drives the acidification of the thylakoid’s lumen through reduction of PQ to PQH2 where the transferred hydrogen ions attenuate LEF by activating qE (Yamori and Shikanai, 2016). In angiosperms, two CEF pathways operate around PSI: a minor NADH dehydrogenase-like (NDH)-complex-dependent and another major pathway involving the chloroplastic protein PGR5-LIKE PHOTOSYNTHETIC PHENOTYPE1 (PGRL1A) and PROTON GRADIENT REGULATION5 (PGR5) (Shikanai, 2007).

PGRL1A is a 28-kDa thylakoid protein with transmembrane domains containing six conserved cysteines facing the stroma whereas PGR5 is a short polypeptide (Hertle et al., 2013). Both *pgrl1ab* and *pgr5* mutant plants exhibited a major decrease (~50%) in the rapid and transient induction of NPQ that occurs during the reactivation of photosynthesis upon exposure to light (Munekage et al., 2002; DalCorso et al., 2008). PGR5 interacts with PGRL1A *in vivo,* and may act as a direct Fd dependent plastoquinone reductase (FQR) (DalCorso et al., 2008; Hertle et al., 2013). PGRL1 controls the trans thylakoid proton gradient and plant growth by stabilizing PGR5 protein (Rühle et al., 2021). Under light fluctuations or stress conditions PGR5 was essential for achieving photoprotection and involved in regulation of donor and acceptor side of PSI (Golding and Johnson, 2003; Yamamoto and Shikanai, 2019). The fine tuning of PGRL1 and PGR5 in CEF regulation is appreciated but the exact mechanism is still unknown as PGR5 has yet to be identified in any purified CEF super complex (Steinbeck et al., 2018).

The CEF and LEF pathways utilize Fd and the activity of the two pathways determines the dynamic redox states of their molecular components (Joliot and Joliot, 2002). Isolated thylakoids showed that chemical reduction with dithiothreitol (DTT) was necessary for high rates of CEF, and suggested that this phenomenon is mediated by a regulatory disulfide intrinsic to PGRL1A (Strand et al., 2015). A regulatory disulfide can switch between the two principal states, the reduced dithiol and the oxidized disulfide, in a reaction that is typically catalyzed by thioredoxins. The reversible interchange between the two states has been shown to control the biological activity of many chloroplast stroma, thylakoid and even luminal proteins (Trebitsh and Danon, 2001; Schürmann and Buchanan, 2008).

The disulfide-regulated chloroplast proteins are generally oxidized in the dark and are reduced upon illumination to activate the phototrophic metabolism during the day (Schürmann and Buchanan, 2008). They are reoxidized during the diurnal transition from day to night and the resultant shift to heterotrophic metabolism. Yet, thioredoxin-controlled oxidizing activity also occurs in the light during the day in a dynamic response to small fluctuations in light intensity. The counterbalance of reducing and oxidizing activities controls the regulatory redox state changes in the light (Trebitsh and Danon, 2001; Martinsuo et al., 2003; Dangoor et al., 2012; Eliyahu et al., 2015; Yoshida et al., 2018). Thus, the activation of CEF via a reducing signal, while beneficial at times such as transitions to light or during light fluctuations, needs to be quenched by re-oxidation during the day for optimal photosynthesis (Strand et al., 2015). Although PGRL1A has been implicated in the control of CEF, the role of its redox regulation in shaping photosynthetic activity during light transitions has not been shown. Here, we examined the redox control of PGRL1A during step changes of steady state low or high light intensity, in fluctuating light and in the presence of inhibitors of photosynthesis. The PGRL1A redox state was determined *in vivo* by measuring the disulfide exchange dynamics and were found to be in concert with modulation of parameters of CEF and NPQ.

## Results

### PGRL1A redox states *in vivo* and *in silico*

Acidic extraction and subsequent treatment of extracts with N-ethylmaleimide (NEM) stabilizes a protein’s redox state. It can then be visualized after fractionation in denaturing but nonreducing fractionation conditions. After night to low light transitions, PGRL1A was previously shown to be trapped into 3 apparent molecular weights of 59, 42 and 28 kDa. The higher molecular weights were sensitive to chemical reduction by dithiothreitol (DTT) and were all reduced to the 28 kDa migration size. Thus the 59 and 42 kDa sizes were assigned as being the oxidized states of PGRL1A and the 28 kDa size represented the fully reduced state (Wolf et al., 2020). The appearance of the reduced 28 kDa form occurred within 10 s during dark to low light transitions (60 μE*m^-2^*s^-1^). However, this change was transient and returned to the more oxidized states after 30 min in constant light (Figure 1, A, B and Supplemental Figure S2D). Importantly, after 30 min under continuous light PGRL1A, once again be reduced in a step transition to higher light (Figure 1B, 600 μE*m^-2^*s^-1^). Note, that no changes in the protein expression level were detected during the experiment, as indicated by similar band intensities obtained under reducing conditions. Thus, the PGRL1A redox state was not only sensitive to the initial dark-light transitions but in a broader sense was reset to encounter further dynamic fluctuations in light; the ramification of which will be examined in this work.

**Figure 1.**
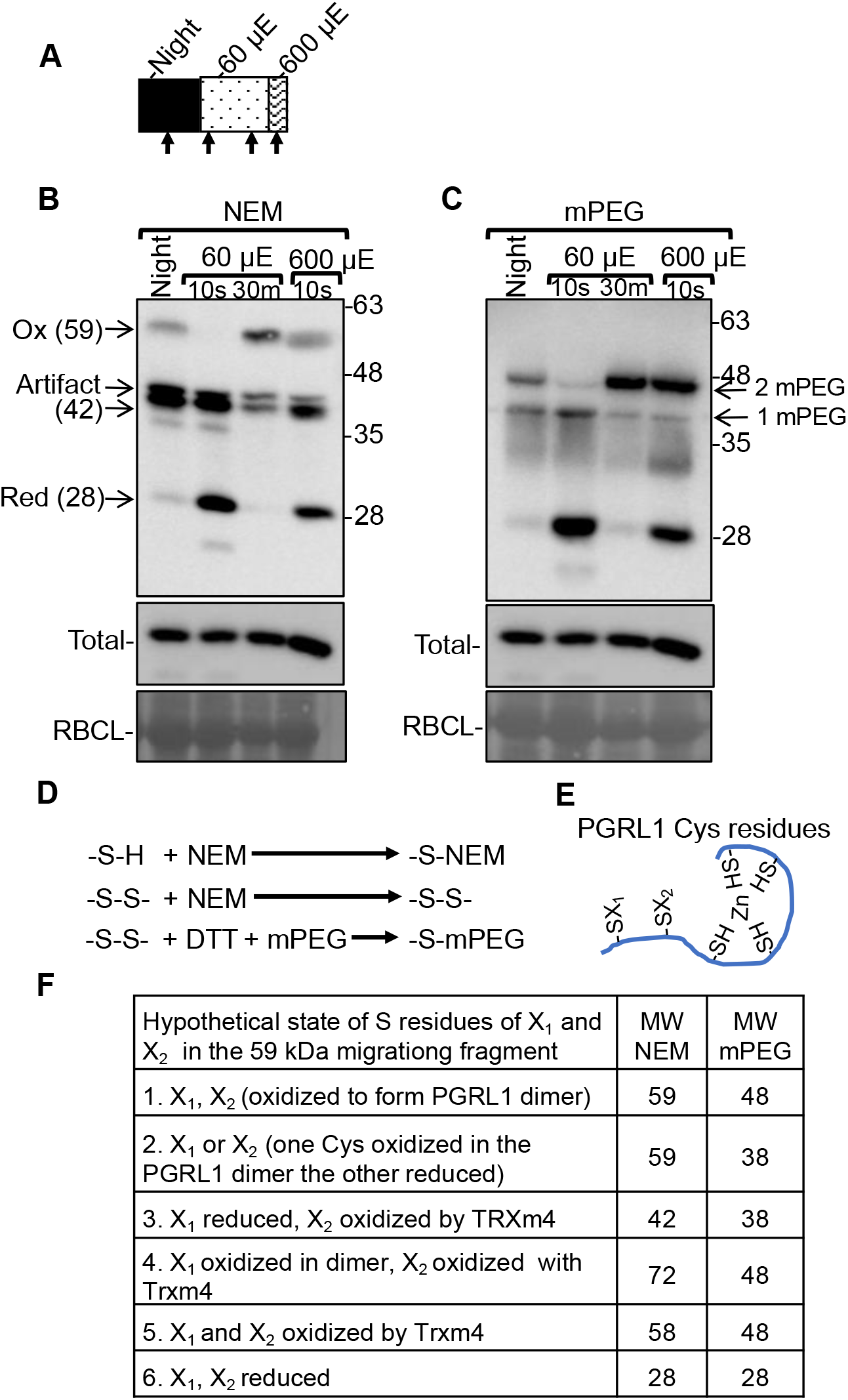
Immunoblot by PGRL1A antibody of NEM and mPEG treated protein extracts after light transitions. A, Schematic representation of light treatments used in (B) and (C) where arrows represent sampling times. B, Immunoblot of night-adapted 4 weeks WT plants labelled with NEM at the end of night and after 10 sec or 30 min light induction at 60 μE*m^-2^*s^-1^ followed by a 10 sec step increase of light intensity to 600 μE*m^-2^*s^-1^. The blots were developed with PGRL1A-specific antibody. Total panel, PGRL1A immunoblot of samples used in the top panel after the addition of DTT. RBCL panel, equal loading control of large subunit of ribulose-1,5-bis-phosphate carboxylase/oxygenase stained by coomassie blue. From top to bottom, arrows point to; the oxidized 59 kDa migration form of PGRL1A; an immunoreactive artifact previously reported in (Wolf et al., 2020); the putative PGRL1A-thioredoxin m4 complex of 42 kDa migration size; the reduced monomeric form of PGRL1A of 28 kDa apparent size. C, Similar to (B) where protein extracts were pretreated with NEM, followed by DTT treatment and subsequently labeled with mPEG. Total protein and RBCL loading controls as in (B). Arrows represent additions of 2 mPEG (48 kDa) or 1 m PEG (38 kDa) to PGRL1A, each mPEG moiety adds approximately 10 kDa to PGRL1A (Rog et al., 2021). The results shown are representative of at least five independent experiments. D, Schematic representation of possible disulfide based redox state of PGRL1A after NEM or NEM followed by mPEG treatments. E, Hypothetical representation of the cysteine residues in PGRL1A as discussed in the text where X1 and X2 correspond to cysteine residues 82 and 183, respectively. F, Table of predicted molecular weights based on NEM or mPEG addition to cysteine residues.

It was of interest to ascertain the molecular characteristics of Cys residues in PGRL1A that could participate in redox interactions. The PGRL1A amino acid sequence contains 6 cysteines and therefore conceivably, PGRL1A could form multiple intramolecular or intermolecular-disulfide bridges (Figure 1, E and F). Recently, it was shown that the 42 kDa thioredoxin m4 formed an intermolecular disulfide with PGRL1A at Cys183 (Okegawa and Motohashi, 2020a). We carried out in-silico structural modeling to examine that possibility and also to deduce additional insights as to the role of the Cys residues. A search against the protein data bank (PDB) using the PGRL1A sequence (324 residues) resulted in no significant similarity to any known protein structure. However, a theoretical structural model for the PGRL1A gene could be created using AlphaFold (Jumper et al., 2021; Varadi et al., 2021). Supplemental Figure S1A shows that while the N-terminal domain of PGRL1A consisting of Met1 to Thr91 containing Cys82 is predicted to be flexible with very low structure confidence (shown within the dotted circle). The C-terminal domain consisting of Ile92 to Ala324 and containing five Cys residues (Cys183, Cys272, Cys275, Cys300 and Cys303 shown in cyan sticks) formed well defined structure.

Comparison with another structure prediction algorithm using the Phyre2 server resulted in a model with 95% confidence, for only part of the C-terminal domain of PGRL1A (ranging from Leu268 to Gln323) containing four Cys residues (Cys272, Cys275, Cys300 and Cys303) (Kelley et al., 2015). Interestingly, this segment of the C-terminal domain was found to be structurally homologous to the zinc binding domain of the Lysine Biosynthetic Protein (LysW) (PDB:3WWN) (Yoshida et al., 2015). Although the sequence identity is only 31%, the four Cys are structurally well aligned suggesting a putative Zn binding site for PGRL1A similar to LysW (Supplemental Figure S1; note the overlap of Cys residues shown in cyan for PGRL1A and in purple for LysW). Structure predication was carried out independently in the trRosetta server (Yang et al., 2020). This resulted in a C-terminal domain (Ile 92 to Ala324) with folding similar to that predicted by the AlphaFold program where the N-terminal domain (Met1 to Thr9) adopts a helical fold (Supplemental Figure S1C). Importantly, the Zn finger motif, part of the C-terminal domain was predicted by all three programs and aligned well with a RMSD of 0.88 Å (Supplemental Figure S1B). However, the deviation between AphaFold and trRosetta predictions of the remaining C-terminal segment (Ile92 to Leu267) is larger with a RMSD of 3.24 Å (area shown within the two dotted circle in Supplemental Figure S1C).

Next, structural theoretical model for the heterodimer of the PGRL1A (without transit peptide) and thioredoxin m4 was created on the AlphaFold (Colab) server (https://colab.research.google.com/github/deepmind/alphafold/blob/main/notebooks/AlphaFold.ipynb). The structural model revealed a heterodimer interface that involves 159 intermolecular interactions (up to 3.5 Å) stabilizing the complex between PGRL1A and thioredoxin m4 (Supplemental Figure S1D; cyan and pink, respectively). These interactions involve 32 residues from the PGRL1A and 30 residues from thioredoxin m4, one of which is S-S bond between Cys183 of PGRL1A and Cys116 of thioredoxin m4 (shown in sticks in Supplemental Figure S1D). No similar stabilized interaction model of Cys82 of PGRL1A with thioredoxin could be established based on the predicted heterodimer structural model. This lends support to the observations of thioredoxin m4 binding made previously (Okegawa and Motohashi, 2020a).

To understand the origin of the ~ 59 kDa migration band, we treated NEM protected proteins with DTT followed by the addition of methoxy polyethylene glycol maleimide (mPEG; Figure 1, C and D). The addition of 1 mPEG moiety to a polypeptide added an apparent ~ 10 kDa. In this case in the dark sample, PGRL1A migrated in two major bands of ~ 48 kDa and ~38 kDa the result of binding of either two or one-mPEG moiety, respectively. In contrast, the 10 sec light exposed sample showed chiefly the 1 mPEG addition (~38 kDa) and the fully reduced 28 kDa form. However, when the 59 kDa migration band was present in the NEM treated extract (samples of night, 30 min at 60 μE*m^-2^*s^-1^ or 10 s at 600 μE*m^-2^*s^-1^; Figure 1B) the 2-mPEG size was always recovered (Figure 1C).

To further establish the origin of the ~ 48 kDa band after mPEG treatment, we extracted the 59 and 42 kDa bands from denaturing nonreducing fractionated NEM treated proteins and proceeded to label those isolated bands with mPEG (Supplemental Figure S3). After mPEG treatment the ~ 59 kDa NEM band was resolved into a major band of ~ 48 kDa which implies that it represents binding of 2 mPEG moieties. In addition, a ~ 38 kDa band appeared which indicates binding 1 of mPEG; possibly the result of partial chemical reactions. In contrast, the ~ 42 kDa NEM band was resolved to ~ 38 kDa, equivalent to one mPEG binding (Supplemental Figure S3). This could likely originate from Cys183 binding to thioredoxin m4 as was reported (Okegawa and Motohashi, 2020). Thus, considering the conserved role of the C-terminal Cys in binding to zinc, it is the Cys82 and Cys183 residues that are likely to be redox sensitive and participate in forming a homo or heterodimers consistent with scenario 1 in Figure 1F. Alternatively, both cysteines could be bound to thioredoxin m4, consistent with scenario 5 in Figure 1F.

### Changes in PGRL1A redox state result in commensurate induction of NPQ and thylakoid membrane potential

We next elaborated on the light sensitive changes of the PGRL1A redox state and their possible physiological significance. Up and down shifts in light regimes were carried out in low and high light as shown schematically in Figure 2, A and F. NEM-treated protein extracts were fractionated on denaturing nonreducing gel systems. Down shifts from 60 to 10 μE*m^-2^*s^-1^ showed no change in the redox state of the prominent 59 kDa oxidized protein (Figure 2B). However, an upshift in light intensity from 10 to 60 μE*m^-2^*s^-1^ showed a transient decrease in the 59 kDa mobility size of oxidized PGRL1A with a concomitant transient increase in the 28 kDa reduced state of PGRL1A (Figure 2B and Supplemental Figure S2). The changes are quantified in Figure 2C. Interestingly, in shifts to high light of 60 to 600 μE*m^-2^*s^-1^ the reduction of PGRL1A was also rapid; yet, was more stable in contrast to that observed in low light transitions (Figure 2, G and H and Supplementary Figure S2). Note, as a loading control parallel treatment were fractionated under reducing conditions (Figure 2, B and G ‘Total’) or stained to reveal the ribulose bisphosphate carboxylase large subunit (Figure 2, B and G; ‘RBCL’).

**Figure 2.**
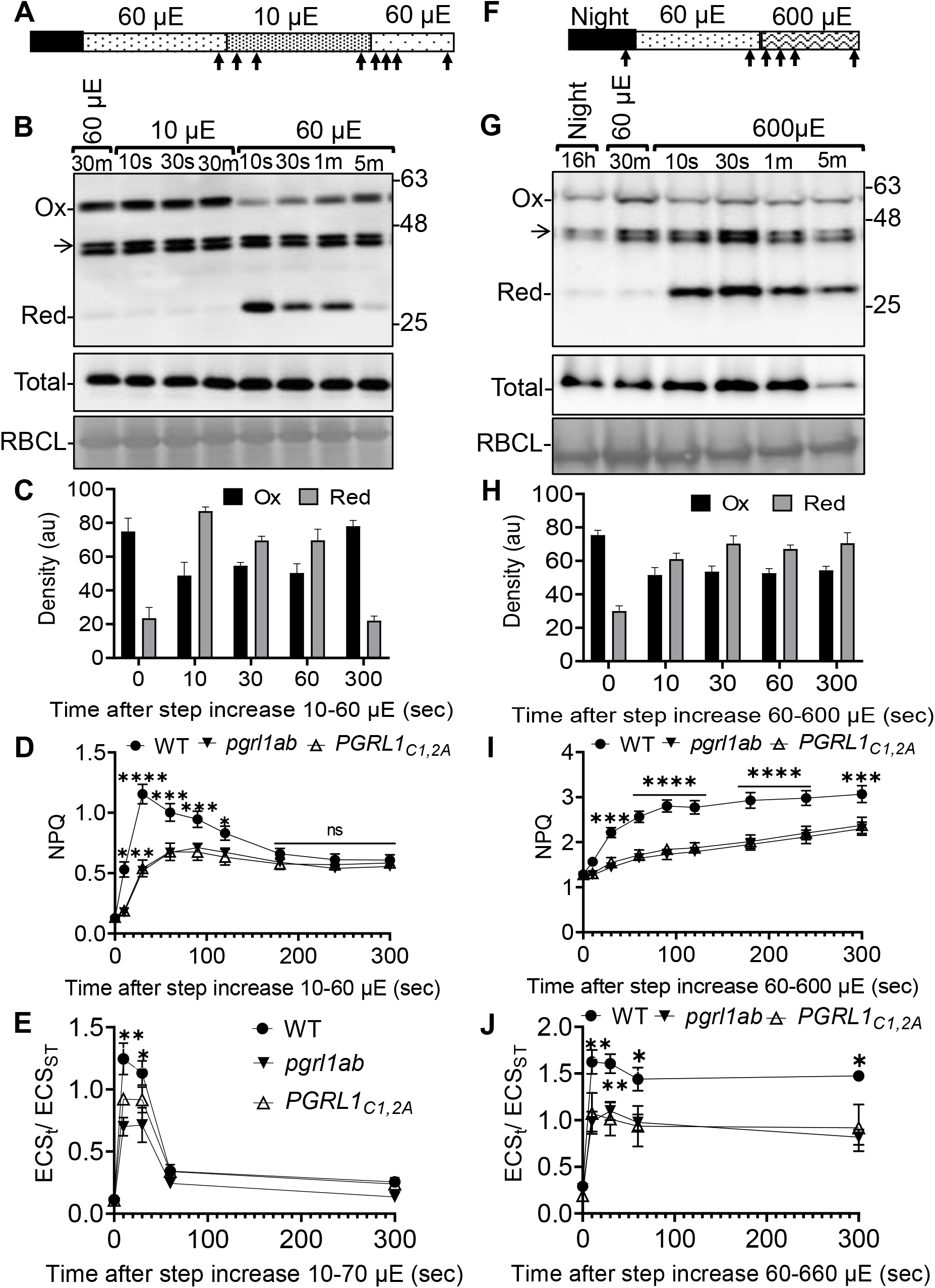
Redox state of PGRL1A changes dynamically during step increases of light intensity. A, Schematic representation of the light regime during step decrease/increase where arrows represent sampling times used in (B). B, Immunoblot showing PGRL1A redox state in WT plants sampled at times and light intensity indicated. Equal loading control panels of ‘Total’ and ‘RBCL’ as in Figure 1. C, Pixel density changes of oxidized and reduced forms of PGRL1A 59 and 28 kDa bands in immunoblot (B) as measured by Image J software. The sampling times are as given in (B) after 10-60 μE*m^-2^*s^-1^ step increase. The pixel density is the average and standard error of 3 biological replicates. D, NPQ induction curve in WT, PGRL1mutants *pgrl1ab* and *PGRL1A_C1,2A_*. Values shown are at step increase in light intensity from 10-60 μE*m^-2^*s^-1^. Each point represents the mean NPQ values of 10 plants with SE at the time points indicated. E, the amplitude of electrochromic shift (ECS_T_/ECS_ST_) of plants at the time points shown in (A). Each data point represents the mean with standard error of 5 biological replicates from 1 leaf of a 21 day old plant. Multiple comparisons were performed by using two-way anova and asterisks indicate a statistically significant difference according to Tukey-Kramer test (*p<0.05, **p<0.01). F-J, Same experimental scheme as in (A-E) at the light intensities indicated. Statistics in D, E, I, and J are statistically significant difference according to Tukey-Kramer test (*p<0.05, **p<0.01, ***p<0.001, ****p<0.0001).

One of the major significances of enhanced cyclic electron flow is the acidification of the lumen and a resultant increase in NPQ (Avenson et al., 2005; DalCorso et al., 2008; Kramer and Evans, 2011). NPQ was measured in wild type (WT) and 2 mutant types of PGRL1A. The mutants are; *pgrl1ab* which does not express PGRL1A and its homolog PGRL1B. Another mutant was PGRL1A_C1,2A_ which is an overexpression mutant form of PGRL1A expressed in the *pgrl1ab* background; where Cys82 and Cys183 were replaced by alanine residues. This inactive polypeptide migrates as if it were a fully reduced form (Wolf et al., 2020). In transitions from 10 to 60 μE*m^-2^*s^-1^ a significantly sharp and transient induction of NPQ was detected in the WT which relaxed slowly to the level displayed by the *pgrl1ab* mutant and mutant of disulfide formation PGRL1A_C1,2A_ (Figure 2D). Significantly, the observed changes follow the time scale of redox state changes in PGRL1A (Figure 2, B and C). The rise in NPQ detected in the mutants was subdued and significantly less then WT. Notable transient changes in proton motive force (PMF) in the WT compared to mutant plants were also revealed by measuring electrochromic shift (ECS_T_). The shifts during the step increase in light intensity of 10-60 μE*m^-2^*s^-1^ were observed to be coherent with the redox changes in PGRL1A (Figure 2E).

The NPQ and ECS_T_ measurements were also found to be in concert with PGRL1A redox state change in transitions from low (60 μE*m^-2^*s^-1^) to high light (600 μE*m^-2^*s^-1^). The WT plants showed significant, but in this case, stable elevated NPQ and elevated ECST values compared to mutant plants (Figure 2, I and J). In all, these finding indicate that the redox changes in PGRL1A are consistent with the regulation of photoprotective mechanisms that are essential for optimizing photosynthetic efficiency. One may deduce that the reduced form of PGRL1A is stimulatory to cyclic flow or reciprocally, that the oxidized form is inhibitory.

### Fluctuating light regimes impact on mutant PGRL1A plants

In order to understand the physiological implications of altered redox homeostasis of PGRL1A and the resultant induction of NPQ; WT and mutant plants, *pgrl1ab* or PGRL1A_C1,2A_ were grown under constant (CL) or fluctuating light (FL). The light regimes were: CL of 10, 60 or 600 μE*m^-2^*s^-1^ during the day under the setting of a day/night (16 h/8 h) cycle. FL was light intensities of 10 μE*m^-2^*s^-1^ for 30 min raised to 60 μE*m^-2^*s^-1^ for 5 min; 60 μE*m^-2^*s^-1^ for 30 min raised to 600 μE*m^-2^*s^-1^ for 5 min and 60 μE*m^-2^*s^-1^ for 5 min raised to 600 μE*m^-2^*s^-1^ (Figure 3). These times were chosen as they were meant to maximize the effect of the temporal difference noted in WT and mutant values of NPQ as deduced from Figure 2D and Figure 2I.

**Figure 3.**
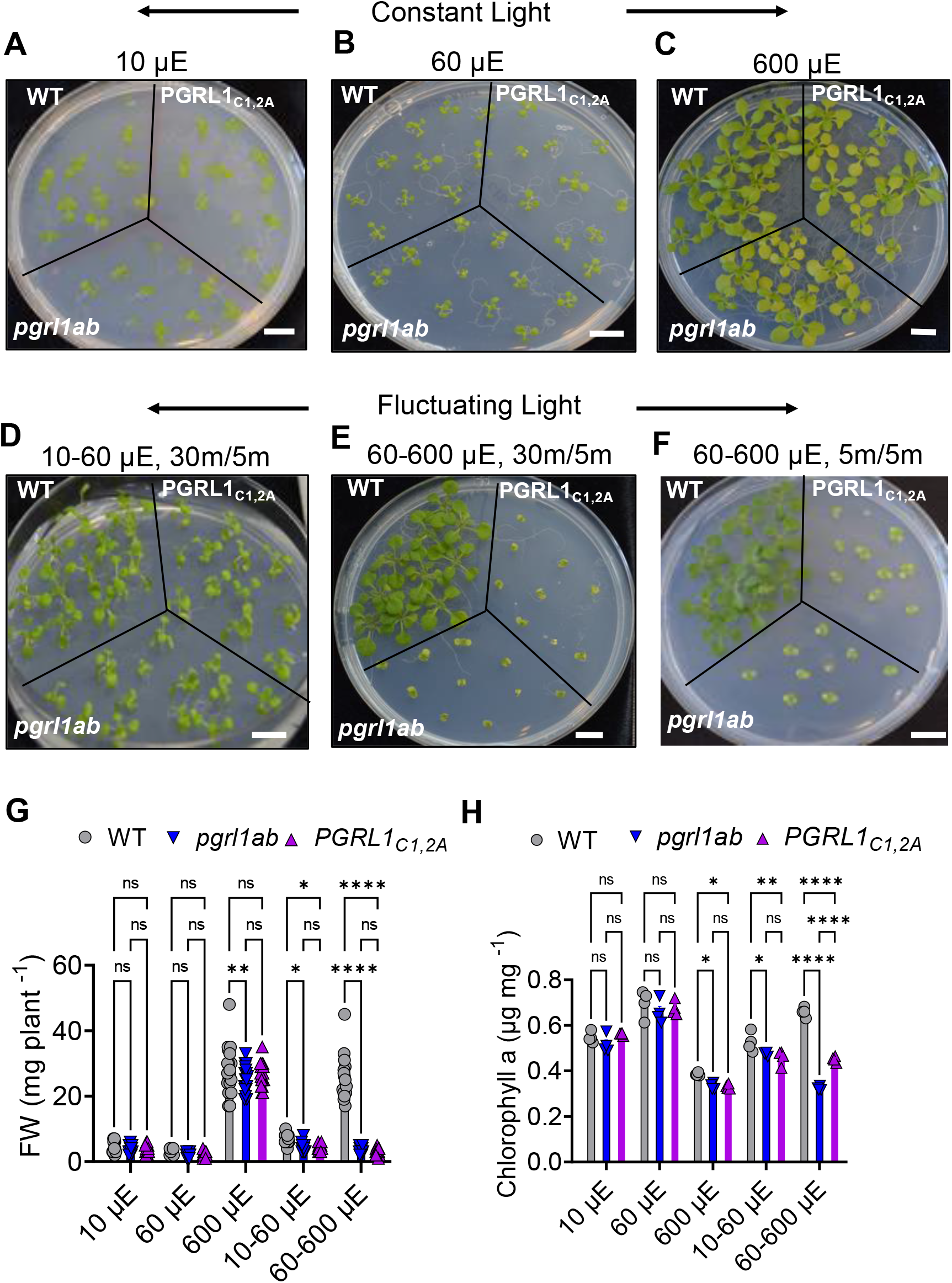
Phenotype variation in WT and mutant plants grown under fluctuating light conditions. WT, PGRL1A mutant *(pgrl1ab)* and plants expressing the mutant form of cysteine *PGRL1A_C1,2A_* were germinated in ½ MS media without sucrose and grown at 22°C in a 16 h constant or fluctuating light to 8 h dark photoperiod. A, constant light of 10 μE*m^-2^*s^-1^. B, constant light of 60 μE*m^-2^*s^-1^ and C, constant light of 600 μE*m^-2^*s^-1^. D, Fluctuating light regimes of 10 μE*m^-2^*s^-1^ (30 min) to 60 μE*m^-2^*s^-1^ (5 min). E, fluctuating light regimes of 60 μE*m^-2^*s^-1^ (30 min) to 600 μE*m^-2^*s^-1^ (5 min). F, fluctuating light regimes of 60 μE*m^-2^*s^-1^ (5 min) to 600 μE*m^-2^*s^-1^ (5 min). G, Fresh weight (mg/plant) of WT, *pgrl1ab* and *PGRL1A_C1,2A_* genotype grown at designated light conditions (A-F) after 2 weeks. The results shown are average of 20 plants. H, Chlorophyll a (μg/mg) of each genotype as in (G) with 3-4 plants/replicate. The results for (G and H) are the average of 3 biological replicates and two-way anova with multiple comparison for each genotype performed with Prism (*v9.*0). Asterisks indicate statistically significant differences according to Tukey-Kramer test (ns, non-significant *p<0.05, **p<0.01, ***p<0.001, ****p<0.0001).

Under CL of 10 or 60 μE*m^-2^*s^-1^, no qualitative or quantitative phenotypic difference was observed as estimated visually (Figure 3, A and B) by fresh weight (FW) or chlorophyll content (Figure 3, G and H). Under CL of high light (600 μE*m^-2^*s^-1^), mutant plants showed slightly impaired growth compared to the WT (Figure 3, C, G and H). When plants were kept under FL regime of 10-60 μE*m^-2^*s^-1^, WT plants showed moderate but significant better performance in terms of higher fresh weight and chlorophyll content compared to mutant plants (Figure 3, D, G and H). Under fluctuating light of 60 to 600 μE*m^-2^*s^-1^as shown in Figure 3, E, F and H, the mutant growth was reduced drastically whether in fluctuating rates of 30 min/5 min or 5 min/5min. The results imply that PGRL1A redox change plays an important role in alleviating stress generated under FL regimes and underscores the significance of NPQ induced by cyclic electron flow in these conditions.

### PGRL1 impacts on the P700 redox state during light transitions

PGR5/PGRL1A mediated flow around PSI is considered to be the major cyclic electron pathway in angiosperms. Under high-light and fluctuating light conditions the PGR5 mutant was shown to provide a protective mechanism for the integrity of PSI (Munekage et al., 2002; Munekage et al., 2004). It was of interest to examine the impact of PGRL1A on the redox state of P700 during step increases in light intensity. Y(ND) represents the ratio of oxidized P700 and is used to monitor the pH-dependent regulation of Cyt b6f complex activity that may occur during cyclic electron flow (Yamamoto and Shikanai, 2019). WT plants showed significantly higher and transient Y(ND) compared to mutant plants at 10 sec or 30 sec after a step increase in light intensity, from 10 to 70 μE*m^-2^*s^-1^. In this case, the PGRL1A state would be largely reduced (e.g. Figure 2); subsequently the Y(ND) values subside. In contrast, mutant lines *pgrl1ab* and *PGRL1A_C1,2A_* barely reacted at 1 min or 5 min after the step increase (Figure 4A). In comparison, after a step increases to higher intensity, 70 to 660 μE*m^-2^*s^-1^ the donor side regulation Y(ND) was stably high at all the time points investigated (10 sec to 5 min) compared to mutant plants, *pgrl1ab* or PGRL1A_C1,2A_ (Figure 4C). Notably, this parallels the extended reduced redox state of PGRL1A (e.g. Figure 2). Reciprocally, Y(NA) represents the ratio of reduced P700 that cannot be oxidized by a saturating flash and is indicative of limitation in electron acceptor activity from PSI (Zhou et al., 2022). During low light transitions Y(NA) values were significantly lower in a transient way in WT plants compared to higher values in mutant plants and they were lower in a stable way after high light transitions (Figure 4, B and D). Thus, Y(ND) and the reciprocal values of Y(NA) both reflect the measured changes in the redox state of PGRL1A.

**Figure 4.**
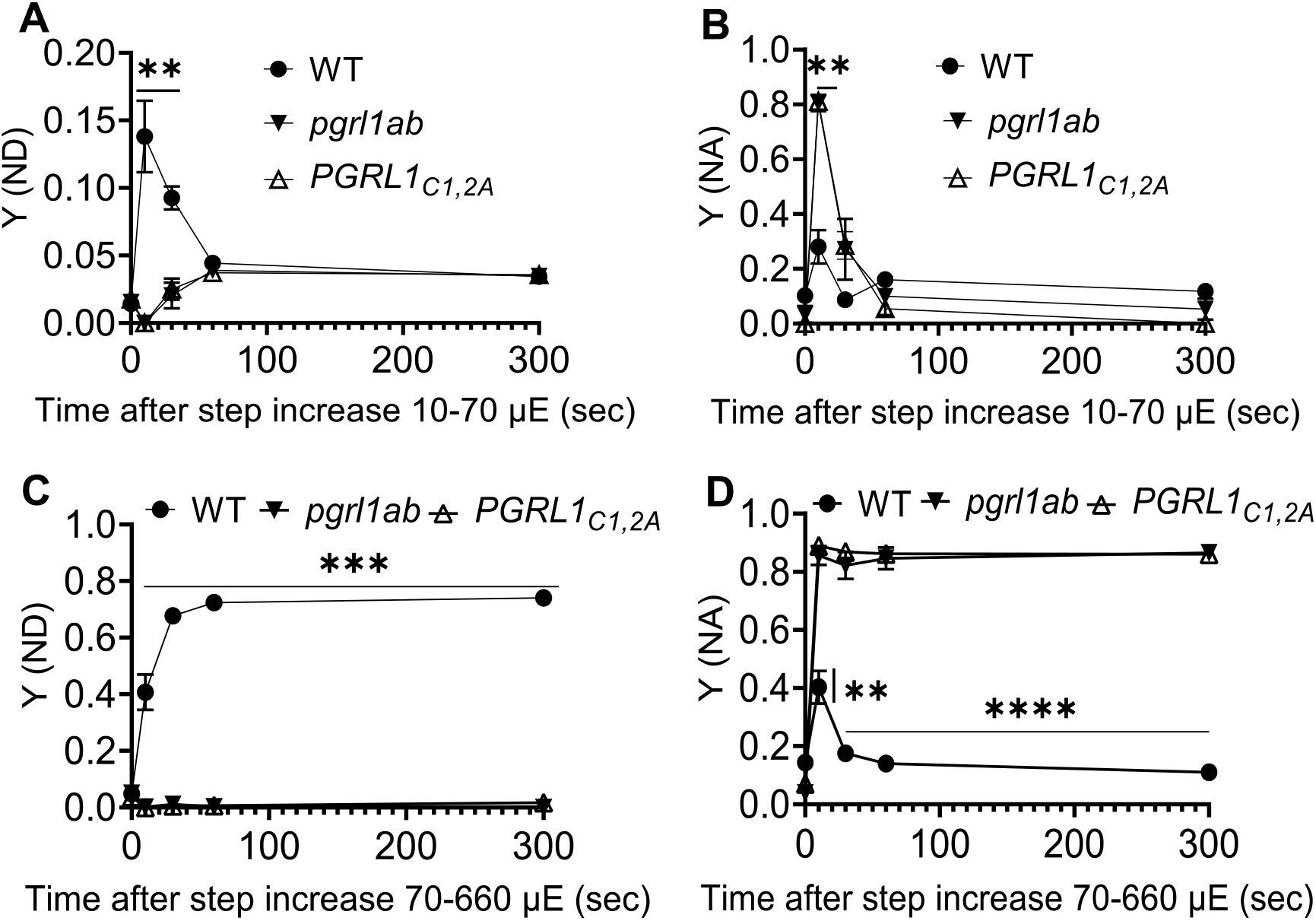
Redox state of PS I after step increases in light intensity. Soil grown 6-week-old WT and *pgrl1ab* and *PGRL1A_C1,2A_* lines were examined for non-photochemical quantum yield of PSI in donor side regulation Y (ND) and acceptor side regulation Y(NA) computed as described in the Materials and Methods during step decrease and increase in light intensity. A and B, Y (ND) and Y (NA) parameters, respectively during step increase at 10-70 μE*m^-2^*s^-1^ light intensity. C and D, Y (ND) and Y (NA) parameters, respectively during step increases of 70-660 μE*m^-2^*s^-1^ light intensity. Each point represents the mean of 5 biological replicates with standard error. Two-way anova with multiple comparison between WT and mutant lines at each time points, were performed with Prism *(v9.0)* and asterisks indicate a statistically significant difference according to Tukey-Kramer test (*p<0.05, **p<0.01, ***p<0.001, ****p<0.0001).

### Redox changes of PGRL1A are modified by inhibitors of photosynthesis

To appreciate the control of the redox state of PGRL1A by photosynthetic electron flow, we investigated the impact of photosynthesis inhibitors. Cyclic electron flow mediated by PGR5/PGRL1A pathways was found sensitive to antimycin A (AA) (Munekage et al., 2002; Munekage et al., 2004; Hertle et al., 2013). Whole seedlings were vacuum infiltrated with AA and the redox state of PGRL1A was examined during dark to light transitions of 60 μE*m^-2^*s^-1^. To determine if the inhibition was effective, the application of AA could be shown to abolish the transient increase in NPQ between WT and PGRL1A mutants (Supplemental Figure S4, A and B). Interestingly, the 59 kDa PGRL1A polypeptide underwent rapid reduction during transition to light and rapidly recovered its oxidized state in a manner similar to WT (Figure 5, A and B). Control ‘total’ PGRL1A immunoreactive polypeptide and stained ‘RBCL’ panel showed that amount of protein did not change significantly during the transition. Based on these observations, although AA directly blocks cyclic electron flow, it is evident that PGRL1A redox state was not affected by AA.

**Figure 5.**
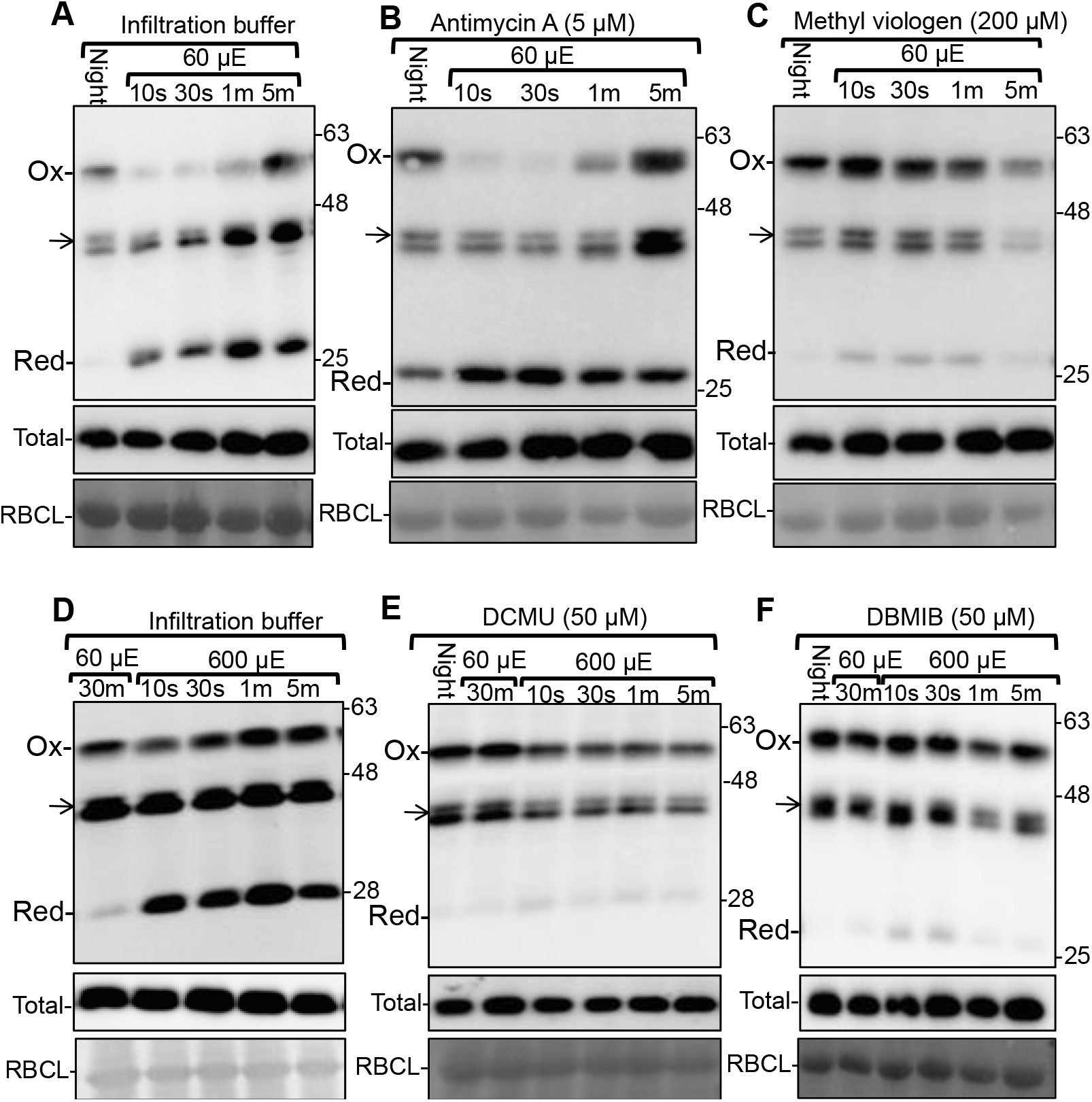
Effect of photosynthetic inhibitors on the redox state of PGRL1A. Immunoblot showing the PGRL1A redox state in WT plants samples with and without inhibitors at times indicated after changes in light intensity. Equal loading controls ‘Total’ and ‘RBCL’ are as in Figure 1. A and D, Plants treated with infiltration buffer under different light regimes as indicated. Treatments are: B, Antimycin A. C, Methyl Viologen. E, DCMU and F, DBMIB.

PGR5/PGRL1A complex participates in the CEF pathway in angiosperms which culminates in the feeding the photosynthetic equivalents to the PQ pool (Munekage et al., 2004; DalCorso et al., 2008; Suorsa et al., 2016). To examine the impact of CEF activity, methyl viologen (MV), a potent electron acceptor downstream of PSI, was employed. The measurement of NPQ during dark to light transitions of 60 μE*m^-2^*s^-1^ showed the expected transient enhanced values compared to water control in WT but was aberrantly elevated after MV treatment (Supplemental Figure S4C). Stable heightened values for NPQ in the presence of MV have been reported previously (Takahashi et al., 2009). Interestingly, PGRL1A was observed to be trapped in its oxidized state of ~ 59 kDa (compare Figure 5, C and A). The ‘total’ and ‘RBCL’ control panels showed that amount of protein did not change significantly during the transition. The results indicate that changes in the acceptor side of PSI as induced by MV impact on changes in the PGRL1A redox state.

The herbicide 3-(3,4-dichlorophenyl)-1,1-dimethylurea (DCMU), binds to the Qa side of PSII and blocks photosynthetic electron transport in the light. In the absence of transport, PQ remains in oxidized state; whereas the quinone analogue 2,5-dibromo-3-methyl-6-isopropylbenzoquinone (DBMIB) occupies the Q_o_ site of the cytochrome b6f complex causing PQ to remain in a reduced state (Metz et al., 1986; Roberts et al., 2004). Plants were examined during a step increase of 60 to 600 μE*m^-2^*s^-1^. As shown in Supplemental Figure S3, E and F, changes in the NPQ states were abrogated in the presence of DCMU and DBMIB, respectively. In all cases, in the presence of either DCMU or DBMIB, PGRL1A was found to be trapped in the oxidized state 59 kDa form when compared to the transient changes in the water control (compare Figure 5, E, F and D). The ‘Total’ and ‘RBCL’ panel showed that amount of protein did not change significantly between the measurements. Based on these results, we conclude that active electron flow is important for reduction of PGRL1A.

## Discussion

Diurnal light fluctuations of even moderate light intensity are considered potent stress factors and an important aspect of CEF induction (Eberhard et al., 2008; Kono and Terashima, 2014; Grieco et al., 2021). Hence, the transient adjustment of photosynthetic rate is critical to allow for a smooth transition between the electron transport and stromal biochemistry during light induction and fluctuations. The activation of CEF was shown to contribute to the induction of qE by stimulating the proton gradient during light induction and during stress conditions (Munekage et al., 2002; Ganeteg et al., 2004; Munekage et al., 2004; Avenson et al., 2005; Hertle et al., 2013; Lee et al., 2019). We showed previously that qE is modulated by a regulatory disulfide of PGRL1A during night to day transitions (Wolf et al., 2020). Here we show that rapid reduction of PGRL1A is concomitant with the induction of qE during step increases to low or high light intensity (Figure 2). This scenario is consistent with the detected transient increased of the proton motive force (PMF) that may be attributed to CEF measured as changes in ECS_T_ (Figure 2). This suggests that reductive activation of PGRL1A will stimulate CEF to accept electrons from the over-reduced Fd in a continuum of light flux contingencies. It also explains why the application of MV inhibitor that leads to increased oxidation of Fd would obviate the reduction of PGRL1A (Figure 5C). MV was shown to diminish CEF activity by competing with Fd for photosynthetic electrons from PSI (Kobayashi and Heber, 1994).

PGRL1A reduced state transiently appears in lower light intensities but remains partially reduced state in high light. The mixed, fully reduced and fully oxidized redox states of PGRL1A were found to be in concert with measured PMF and the resultant NPQ (Figure 2). This suggests that reduction of PGRL1A activates the CEF for short or long-term acclimation responses (DalCorso et al., 2008; Kramer and Evans, 2011; Kono and Terashima, 2014; Strand et al., 2016). As CEF contributes to balancing ATP/NADPH ratio, it may indicate the need to meet different physiological demands at different light intensities (Allen, 2002). The implication is that CEF is part of ongoing physiology in high light but is only temporally necessary in low light. In this respect it has been proposed that the alternative non-PGRL1/PGR5-dependent CEF route offered by NADH dehydrogenase-like (NDH) complex operates especially in low light in rice (Yamori et al., 2015). The transience of PGRL1A reduction in low light would also mean that mechanisms to bring about its oxidation are saturated at higher light. By what mechanism could PGRL1A be oxidized? PQ was shown to generate a redox signal to reflect the fluctuations in photosynthetic electron transport (PET) induced by light intensity and regulate the ratio of PSII and PSI (Pfannschmidt et al., 1999). However, PQ was not shown to be involved in disulfide exchange reactions. Alternatively, the PGRL1 redox state could be in equilibrium with Fd and thioredoxins. Indeed, PGRL1was identified in CEF super complex along with other PSI components like Fd in *Chlamydomonas* by immunoblot and mass spectrometry analysis (Steinbeck et al., 2018).

The reformation of the 59-kDa oxidized protein would result in diminishing levels of CEF and in turn relaxation of the trans-thylakoidal proton gradient resulting in quenching of NPQ and CEF (Figure 2). It would appear from the experiments here, that it is the reduced state of PGRL1A that stimulates the trans-thylakoidal proton gradient. This in turn, would induce qE via changes in the PsbS aggregation state and restrict LEF (Li et al., 2000; Tikkanen et al., 2011; Nikkanen and Rintamäki, 2019). For example, it is the reduced state of a redox regulated sarco/endoplasmic reticulum calcium transport ATPase (SERCA2b) that is the active state. The protein was inactivated by oxidation of luminal cysteine pair but regained functionality upon reduction (Ellgaard et al., 2018).

To discern the nature of the six conserved cysteines and ~ 59 kDa migration size, we employed modelling complimented by redox sensitive NEM and mPEG methodology. Any model for PGRL1 dynamic structure needs to consider that 2 Cys residues are oxidized in the 59-kDa complex (Figure 1C and Supplemental Figure S3). The higher molecular weight may be consistent with the formation of a homodimer or binding of 2 molecules of thioredoxin m4 (Figure 1). The redox additions of NEM and mPEG are consistent with both scenarios. In support of homodimer formation, recombinant PGRL1A was reported to form homodimers *in vitro* (Hertle et al., 2013). Furthermore, although one cysteine Cys 123, was shown to be engaged with thioredoxin m4 by disulfide exchange *in vivo,* the other, Cys82, could not be modelled here and was not detected *in vivo* (Okegawa and Motohashi, 2020). It is noted that a homodimer model would require the removal of thioredoxin m4 to be consistent with the observed molecular weights.

Structural modelling by Alphafold, trRosetta, and Phyre2 algorithms (Supplemental Figure S1) provided the unexpected finding that C-terminal domain of PGRL1A containing four Cys residues (Cys272, Cys275, Cys300) exhibited structural homology to the Zinc binding domain of LysW that contains a TFIIB-type zinc finger (Zhu et al., 1996). The function of zinc fingers is to promote protein-nucleic and protein-protein interactions and stabilize proteins. The presence of zinc fingers may explain why mutation of any of the 6 Cys culminated in an inactive protein (Wolf et al., 2020). The zinc finger in PGRL1A may also act as a site for interaction with a regulatory subunit. One such interacting protein candidate could be PGR5 which was shown to interact with the C-terminal stromal exposed portion of PGR1A (DalCorso et al., 2008). It is of interest that this C-terminal portion was also shown, by *in vitro* assay with recombinant protein, to be associated with Fe binding in a redox-sensitive manner (Hertle et al., 2013). It is known that in the absence of the preferred cofactor, zinc fingers will bind Fe (Conte et al., 1996). Another chloroplast protein, the FtsH5 Interacting Protein, was found to contain zinc fingers that facilitated binding to the thylakoid FtsH proteases (Borsani et al., 2018). The latter complex together with Deg proteases, is involved in degradation of photodamaged PSII D1 protein. It would be of future interest to examine for this possible turn-over functionality in PGRL1A as well.

PSI may act as the efficient quencher of excess excitation energy and was shown to be less susceptible to photodamage compared to PSII (Tikkanen and Aro, 2014; Jahns et al., 2017). This quality is due to the modulation of CEF that controls electron flow from PSII to PSI (Munekage et al., 2002; Pascal et al., 2005; Eberhard et al., 2008). Here, we show that mutant plant *pgrl1ab* and *PGRL1A_C1,2A_* displayed higher Y (NA) i.e. had more reduced P700 compared to WT during step increases in light intensity (Figure 4, B and D). Conversely, more effective photosynthetic connectivity was displayed by WT plants in higher donor side values of Y (ND) in WT compared to mutants during step increases in light intensity (Figure 4, A and C). These photosynthetic controls of donor and acceptor sides are hallmarks of protection of PSI and followed the time scale of redox change of PGRL1A; displaying transient or stable control at step increases in light intensity from 10-60 μE*m^-2^*s^-1^ or 60-600 μE*m^-2^*s^-1^, respectively. PGR5 mutants were also shown to have impaired ability to oxidize P700 (Suorsa et al., 2012). Recently, it was found that artificially enhanced donor side and acceptor side regulation were essential for PSI protection under fluctuating light conditions (Jahns et al., 2017; Yamamoto and Shikanai, 2019; Zhou et al., 2022). Together, these findings suggest that PGRL1A redox state regulates CEF activity and is thus involved in the photoprotection of PSI.

Artificial light fluctuations regime was imposed on WT and mutant lines to understand the need for PGRL1A redox changes in step increases in light intensities from 10-60 μE*m^-2^*s^-1^ or from 60-600 μE*m^-2^*s^-1^ (Figure 3). Severely impaired growth phenotype of mutant compared to WT plants were observed and may be attributed to photoinhibition due to an imbalance in the redox poise failing to regulate qE and therefore unable to exert effective photosynthetic controls. The results are similar to that found for the role of CEF under fluctuating light condition in the PGR5 mutant (Suorsa et al., 2012; Kono et al., 2014; Kono and Terashima, 2014; Suorsa et al., 2016).

Inhibitors of photosynthesis were employed to understand the impact of PET on the redox regulation of PGRL1A. CEF was previously shown to be sensitive to AA (Munekage et al., 2002; DalCorso et al., 2008; Hertle et al., 2013; Sugimoto et al., 2013). PGRL1 is not likely the target for CEF inhibition by AA but rather its interacting partner, PGR5 (Rühle et al., 2021). Here, the addition of AA did not prevent the light-induced reduction of PGRL1A *in vivo*, however the transient NPQ induction subsided to the level of mutant plants *pgrl1ab* or *PGRLIA*_C1, 2A_ (Figure 5, A and B; Supplemental Figure S4, A and B). Thus, while changes in the redox state of PGRL1A are essential for regulating NPQ they are not sufficient.

The photosynthetic inhibitors DCMU or DBMIB block the photosynthetic electron transport but result in either oxidized or reduced PQ, respectively. None the less, PGRL1A was found to maintain the oxidized state during all step increases in light intensity (Figure 5, D and E). However, a strong reduction of NPQ value were observed in seedlings infiltrated with DCMU or DBMIB compared to control (buffer control) (Supplemental Figure S4, C and E, F). A similar observation (lower NPQ) was observed in dinoflagellate *Symbiodinium* with DCMU after 5 min light illumination (Lepetit et al., 2013; Aihara et al., 2016). Hence under these conditions, the redox state of PQ is not directly linked to the PGRL1A redox state. These results together with the application of MV imply that the redox state of Fd is involved in controlling reduction of PGRL1A. Our results show that redox state changes in PGRL1A are keyed into the rates of electron flow providing crucial optimization of photosynthesis during light induction or light fluctuation.

## Materials and Methods

### Plant material, growth conditions and chlorophyll analysis

*Arabidopsis thaliana var.* Columbia (Col-0) (WT) was grown in ambient air on solid half-strength Murashige and Skoog (MS) medium in 0.8% agar plates. Plants for immunoblotting were grown under a 16/8 h light/dark cycle at 20°C, at a light intensity of 60 μE*m^-2^*s^-1^ for 3-4 weeks. For phenotyping plants under fluctuating regimes, WT, *pgrl1ab* or *PGRL1A_C1,2A_* were grown in a customized growth chambers where light during the days were fluctuating at 10 or 60 μE*m^-2^*s^-1^ for 30 min or 5 min, respectively. For higher light conditions, plants were grown in light fluctuations of 60 or 600 μE*m^-2^*s^-1^ for 30/5 min or 5 min, respectively. Growth was at 8/16 h light/dark cycle at 20°C for 2 weeks before documenting the phenotype. The double mutant homozygous line (*pgrl1ab*) and *PGRL1A_C1,2A_* genotype have been described (Wolf et al., 2020). For chlorophyll analysis, samples of 2-week-old seedlings were homogenized in 1 ml methanol (−20°C) and centrifuged for 5 min at 13,000g. All steps were performed in the dark. Calculations for chlorophyll content were as described in (Porra et al., 1989).

### Structural theoretical modeling

The structural model for the PGRL1A compared three different methods. The AlphaFold that uses an artificial intelligence (AI) algorithm to predict highly accurate protein structure. The trRosetta that uses neural network for fast and accurate structural prediction (Jumper et al., 2021; Varadi et al., 2022) (https://yanglab.nankai.edu.cn/trRosetta) (Yang et al., 2020) The Phyre2 protein homology analogy recognition server http://www.sbg.bio.ic.ac.uk/phyre2/html/page.cgi?id=index (Kelley et al., 2015). The structural prediction for the heterodimer of the PGRL1A (without transit peptide) and the thioredoxin m4 out used the AlphaFold (Colab) server (https://colab.research.google.com/github/deepmind/alphafold/blob/main/notebooks/AlphaFold.ipynb).

### Protein disulfides assay and Immunoblot Analyses

For determining redox states of the PGRL1A protein, plant extracts containing 4-5 seedlings/sample were immediately harvested by grinding in 10 % trichloroacetic acid (TCA) followed by centrifugation to obtain the protein pellets. The acidic conditions protonate Cys residues preventing further thiol exchanges. Protein extraction and redox labeling were carried out as previously described (Dangoor et al., 2012) except that the resuspension of protein precipitate was in modified denaturing urea buffer (8 M urea, 2% SDS, 100 mM HEPES-KOH pH 7.4, 10 mM EDTA and protease inhibitor [Calbiochem, now Merck, https://www.merckmillipore.com] containing 50 mM N-ethylmaleimide (NEM) for 30 min. To label the oxidized cysteines with mPEG, the extracts were further reduced by 100 mM DTT, excess DTT was removed by 100% acetone precipitation resuspend in modified denaturing urea buffer containing 2 mM mPEG and fractionated by reducing or nonreducing SDS/PAGE, as indicated (Rog et al., 2021). Proteins were analyzed by immunoblotting with PGRL1A-specific polyclonal antibody raised against the N-terminal loop (aa 61 to 200; N-PGRL1A) of PGRL1AA in rabbits.

### Chlorophyll fluorescence analysis and P700 absorbance change

Plants were grown in ambient air on solid ½ MS medium with 0.8% agar under 8/16 h light/dark cycle at 80 μE*m^-2^*s^-1^ at 20°C/18°C, respectively. Chlorophyll fluorescence was measured in 21day old (n=10 per measurement), using an Imaging PAM chlorophyll fluorometer (Heinz Walz GmbH). Actinic light of 60 μE*m^-2^*s^-1^ was switched on 40 sec after F0 and Fm determination. Saturating pulses (SP) of 10,000 μE*m^-2^*s^-1^ were administered for 600 ms, at designated intervals in a customized script during light transitions. NPQ was calculated as [(Fm-Fm’) /Fm’] where Fm represents the maximum fluorescence emission recorded in dark-adapted leaves, and Fm’ represents the maximum fluorescence value recorded at specific time intervals during the illumination phase.

Chlorophyll P700 absorption changes of the PSI reaction center were measured using DUAL-PAM-100 measuring system for simultaneous assessment of P700 and chlorophyll fluorescence analyzer equipped with a P700 dual-wavelength emitter at 830 and 870 nm; Walz). The redox changes of P700 was assessed in attached leaves of 6 weeks soil grown *Arabidopsis* genotypes in night adapted plants or 1-hour dark incubation were used prior to measurement. Pm was determined by the application of an SP in the presence of far-red (FR) light. The maximal level of oxidized P700 during AL illumination (Pm’) was determined by application of SP. Y(NA) was calculated as (Pm-Pm’)/ Pm and Y(ND) was calculated as (P/Pm).

### Electrochromic shift analysis

Plants grown on soil under 8/16 h light/dark cycle at 60 μE*m^-2^*s^-1^ at 20°C/18°C, respectively for 6-weeks prior the experiments. Attached leaves of night-adapted or 1h dark incubated plants were used for measuring ECS. The ECS was monitored as the deconvoluted absorbance change at 515 nm by using a DUAL-PAM-100 (Walz, Germany) equipped with a P515/535 emitter-detector module. After actinic light of 10, 60 or 600 μE*m^-2^*s^-1^ was switched on, it was switched off at designated time points and ECS_T_, which represent the difference in total PMF between light and dark, was calculated from the total amplitude of the rapid decay of the ECS signal approximately 0.3 sec after the switch to the dark as described in (Avenson et al., 2005). The ECS_T_ signals were normalized against the 515-nm absorbance change induced by a single turnover flash (ECS_ST_) as measured in dark adapted leaves before recording light induced change in ECS to circumvent the changes in leaf thickness and chloroplast density among leaves.

### Photosynthetic inhibitors treatment

Seedlings (3-4 weeks old) were vacuum infiltrated with medium containing 300 mM sorbitol, 20 mM HEPES/KOH (pH7.5), 5 mM MgCl2 and 2.5 mM EDTA with 5 μM antimycin A, or 50 μM DCMU or DBMIB or 200 μM MV for 5 min (Sugimoto et al., 2013). After vacuum infiltration, extra liquid was removed after placing seedlings on dry blotting sheets, placed on half strength MS plates and kept in dark for 30 min before fluorescence or light treatment for immunoblotting.

## Acknowledgements

This paper is dedicated to the memory of Prof. Avihai Danon as an honored scientist and mentor who made seminal contributions to our understanding of redox regulation. We thank Dr. Shilo Rosenwasser for his critical comments on the manuscript. This research was supported by grants from the Israel Science Foundation; Grants No. 1013/17 and 2106/21.

## Author Contributions

A.K.C.: conceptualization, methodology, investigation, software, formal analysis, and writing-original draft, review and editing. O.D.: structural theoretical modelling, writing-review and editing. R.F.: conceptualization, supervision, writing-original draft, review and editing.

## Data Availability Statement

Data supporting the findings of this work are provided in the main text and the supporting information files. All data and materials used in this study will be available from the corresponding author.

## Figure legend of the supplemental figures

**Supplemental Figure S1:** Ribbon representation of the PGRL1a predicted structure.

**Supplemental Figure S2.** The redox state of PGRL1A changes dynamically during diurnal and step changes in light intensity

**Supplemental Figure S3:** Immunoblot of excised bands that were treated with DTT and mPEG.

**Supplemental Figure S4:** NPQ levels in vacuum infiltrated plants with photosynthetic inhibitors

## Abbreviations

NPQ: nonphotochemical quenching
PGR5: PROTON GRADIENT REGULATION5
PGRL1A: PGR5-like phenotype 1A
DTT: dithiothreitol
NEM: N-ethylmaleimide
CEF: cyclic electron transport
LEF: linear electron transport

## Supplemental Figure S1-S4

**Supplemental Figure S1.**
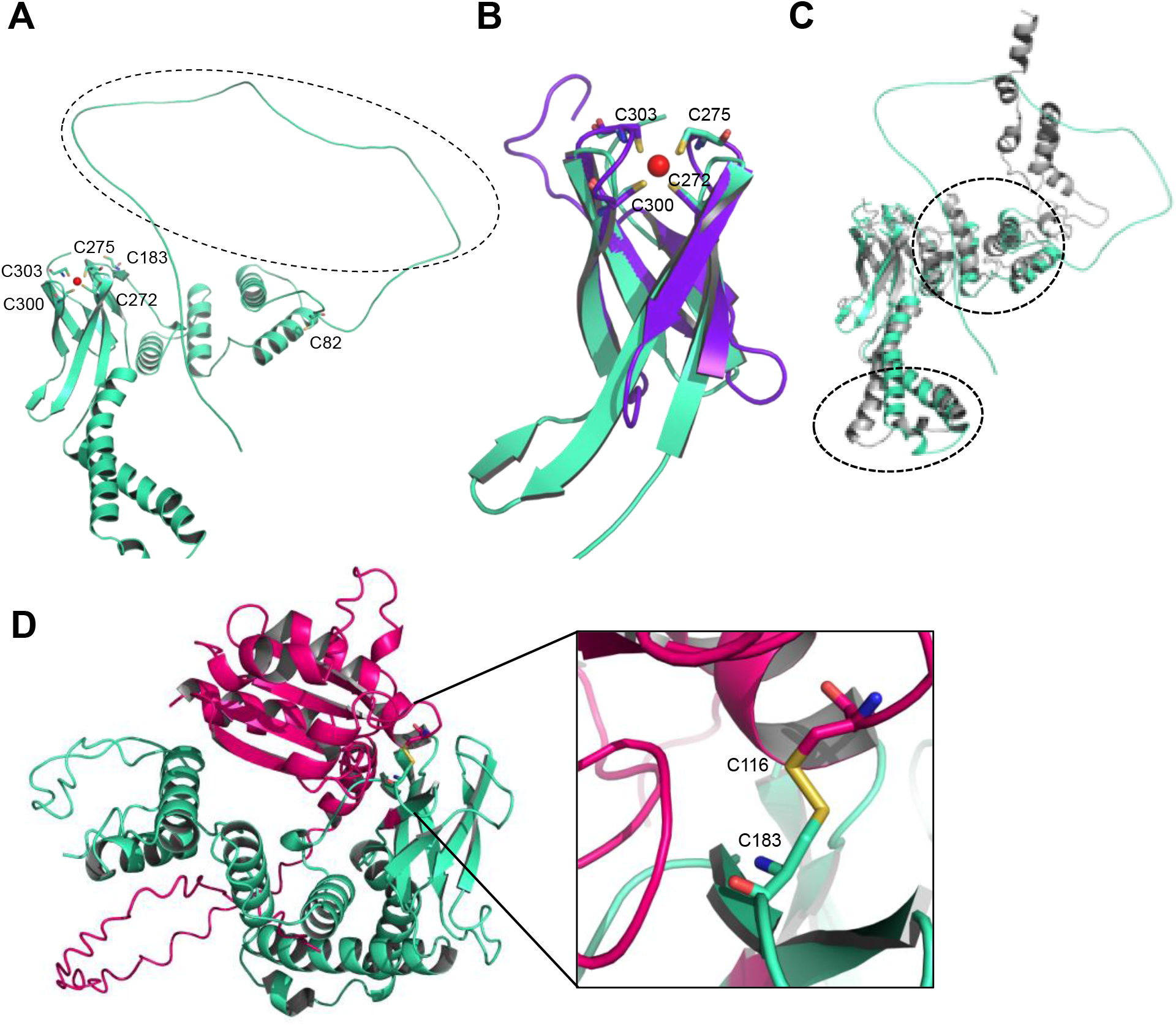
Ribbon representation of the PGRL1A predicted structure. A, AlphaFold PGRL1A predicted structure. The area of the flexible N-terminal domain consists of residues Met1 to Thr91 is shown in dashed black circle. The six Cys residues (Cys82, Cys183, Cys272, Cys275, Cys300 and Cys303) are in stick representations. B, The region ranging from Leu268 to Ala323 of PGRL1A is predicted to be structurally related to the Zinc binding domain of LysW (PDB:3WWN). The four Cys residues (Cys272, Cys275, Cys300 and Cys303) in the putative Zn binding (red colored ball) site of PGRL1A are shown by stick representation (cyan colored). The LysW zinc finger structure is in purple. C, Alignment of PGRL1A predicted structure utilizing AlphaFold and TrRosetta (cyan and grey respectively). D, AlphaFold predicted structure of the PGRL1A and the Thioredoxin m4 heterodimer (cyan and pink respectively). The predicted S-S bond between Cys116 of Thioredoxin m4 and Cys183 of PGRL1A are shown in sticks. Figures were created using PyMOL.

**Supplemental Figure S2.**
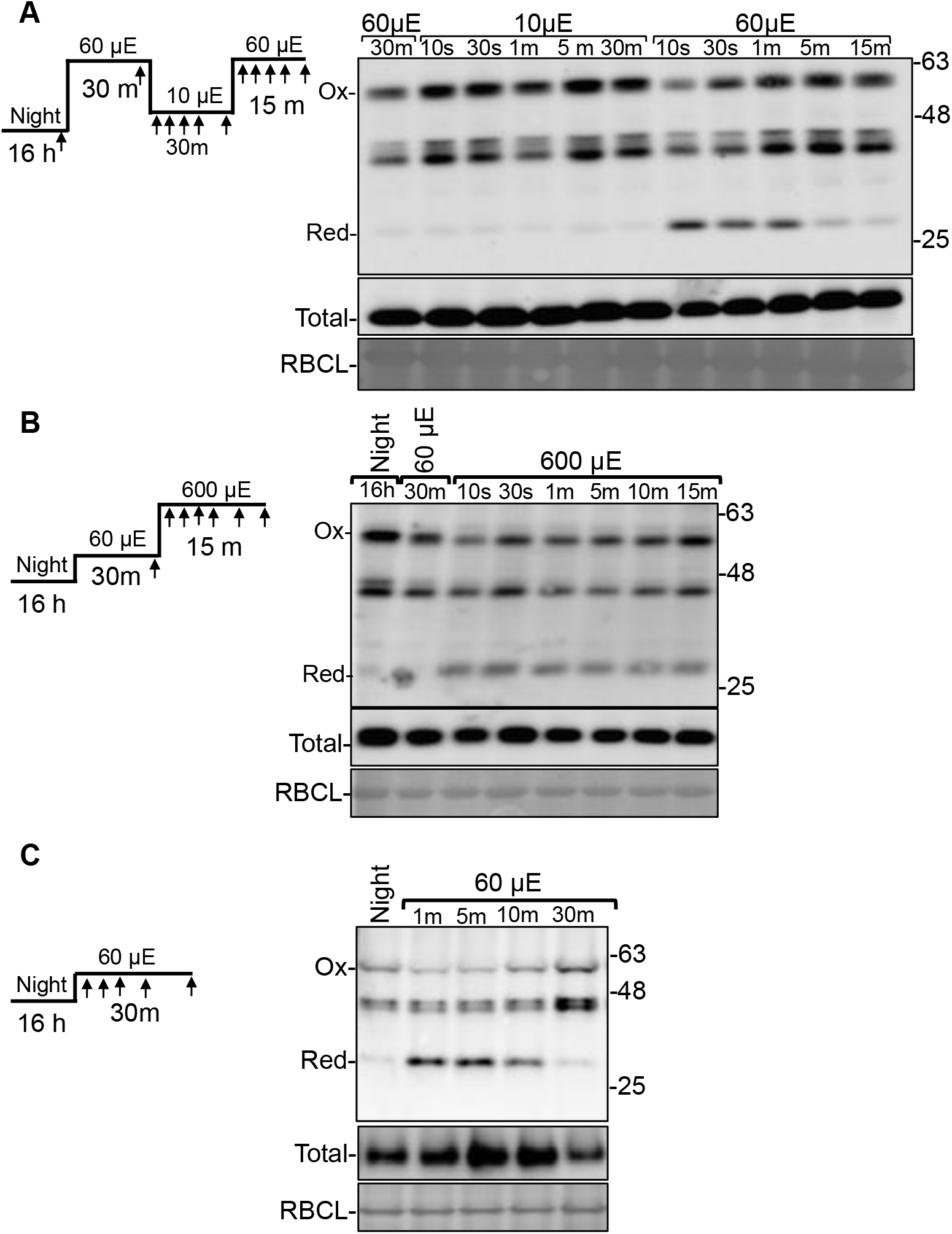
The redox state of PGRL1A changes dynamically during diurnal and step changes in light intensity. A, On the left, the schematic representation of the low light regimes during step decrease/increase where arrows represent sampling times as shown in the immunoblot on the right. B, as in (A) at higher light intensities. C, as in (A) for night to low light transition. Equal loading controls: Total panel, immunoblot of the PGRL1 immunoreactive proteins in top panel fractionated under reducing conditions. RBCL panel, large subunit of ribulose-1,5-bis-phosphate carboxylase/oxygenase stained by coomassie blue. The oxidized 59 kDa and the reduced monomeric form of PGRL1A of 28 kDa apparent size are indicated.

**Supplemental Figure S3.**
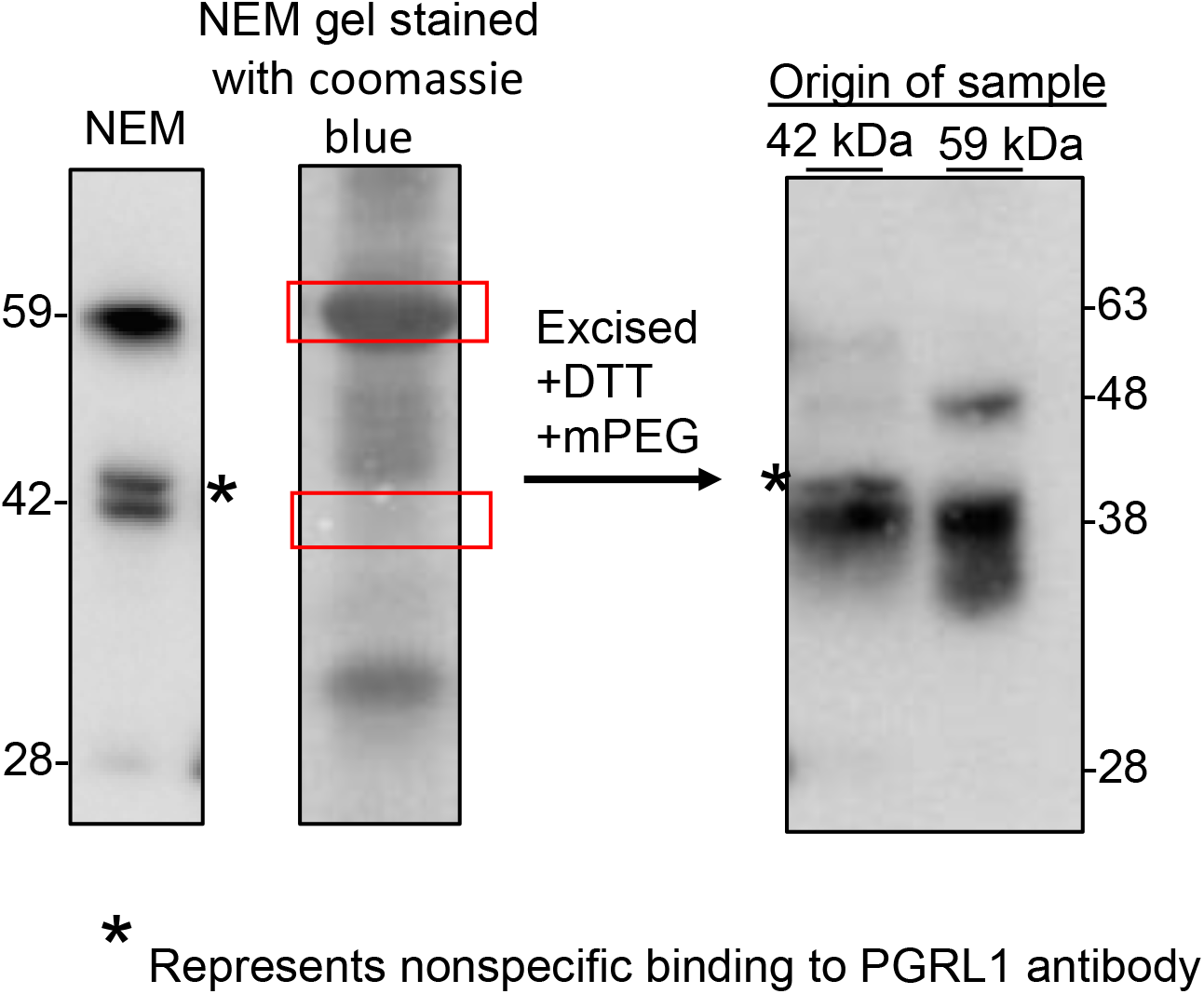
Immunoblot of excised bands that were treated with DTT and mPEG. Immunoblot of night-adapted 4 weeks WT plants labelled with NEM at the end of night. The blots were developed with PGRL1A-specific antibody (gel on left) or stained with coomassie blue (CBB; gel in center). The 59 and 28 kDa regions were excised and treated with DTT and mPEG as described in the Materials and Methods and fractionated on a nonreducing denaturing gel and processed with PGRL1A antibody. Asterisk marks non-specific band.

**Supplemental Figure S4.**
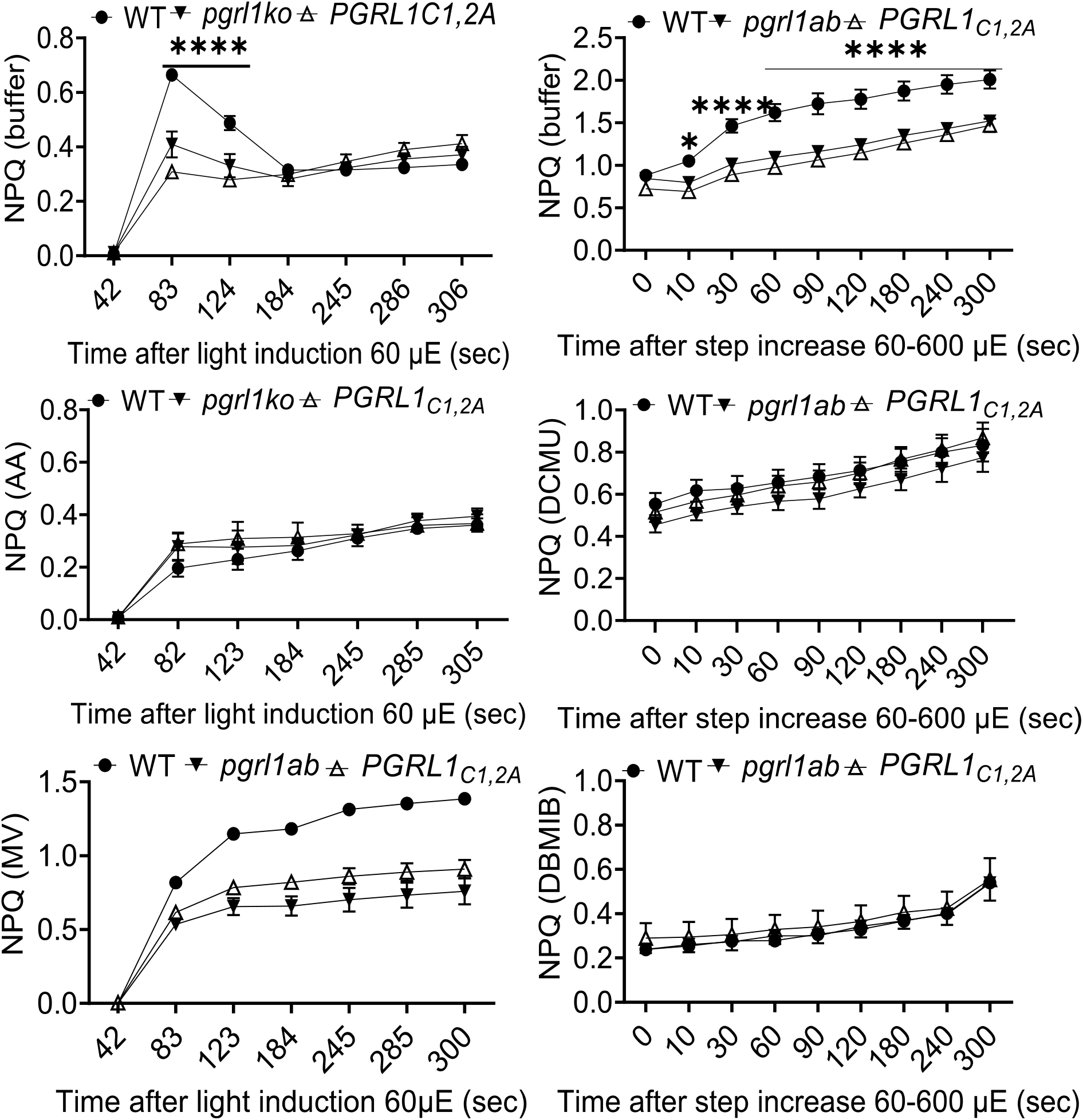
NPQ levels in vacuum infiltrated plants with photosynthetic inhibitors. NPQ was measured in WT and mutant genotype *pgrl1ab* and PGRL1_C1,2A_ seedlings after vacuum infiltration for 5 min with various photosynthetic inhibitors. A-C, dark to light (60 μE*m^-2^*s^-1^) induction. D-F, step increase in light intensity at 60-600 μE*m^-2^*s^-1^. Measurements in (A and D) are of control infiltration buffer without inhibitors. The inhibitors used are; B, Antimycin A (AA); C, Methyl viologen (MV); E, DCMU and F, DBMIB.

